# Dopamine transporter threonine-53 phosphorylation dictates kappa opioid receptor mediated locomotor suppression and conditioned place aversion via transporter upregulation

**DOI:** 10.1101/2024.05.09.593368

**Authors:** Durairaj Ragu Varman, Lankupalle D. Jayanthi, Sammanda Ramamoorthy

**Author notes:** Corresponding author: Sammanda Ramamoorthy, Department of Pharmacology and Toxicology, Virginia Commonwealth University, Richmond, VA 23298.; or.

## Abstract

Dynorphin (DYN)/kappa opioid receptor (KOR) activation contributes to aversion, dysphoria, sedation, depression, and enhanced psychostimulant-rewarding effects, which have been attributed to the inhibition of dopamine (DA) release. DYN fibers synapse onto DA terminals which express both KOR and dopamine transporter (DAT). DAT activity is critical in the regulation of DA dynamics and dopaminergic neurotransmission. Previously, we demonstrated that KOR agonists upregulate DAT activity via ERK1/2 signaling involving phospho-Thr53 DAT (pT53-DAT). However, whether pT53-DAT is involved in KOR-mediated DAT regulation in-vivo and whether such phenomenon contributes to the behavioral effects of KOR agonism are unknown. Here, we investigated the role of endogenous pT53-DAT in KOR-mediated DAT regulation and the effect of KOR agonists on locomotor suppression and aversive behaviors using DAT-Ala53 knock-in mice expressing DAT carrying non-phosphorylatable Ala at position 53 replacing Thr. Acute systemic administration of KOR agonist, U69593 resulted in KOR antagonist-sensitive increases in DAT activity in parallel to increases in pT53-DAT, and DAT V_max_ and surface expression in the ventral and dorsal striatum (containing the nucleus accumbens and caudate putamen respectively) of WT, but not DAT-Ala53 mice. KOR agonists produced conditioned place aversion (CPA) and locomotor suppression in WT but not DAT-Ala53 mice. However, both WT and DAT-Ala53 mice exhibited similar lithium chloride-induced CPA and morphine-induced conditioned place preference (CPP). These findings provide the first evidence that locomotor suppression and aversive responses to KOR agonists manifest due to the modulation of DAT activity via DAT-T53 phosphorylation establishing a causal relationship of pT53-DAT in KOR-mediated DAT regulation and KOR agonist-induced adverse effects.

## Introduction

The dynorphin (DYN) and its native kappa opioid receptor (KOR) are expressed in the central nervous system and play a role in many physiological functions [1–4]. KOR activation has been implicated in dysphoric and psychotomimetic effects in humans [5] and aversive behaviors in rodents [6–8]. Systemic administration of KOR agonists, including U69593, U50488 and salvinorin A (SalA) induces conditioned place aversion (CPA) and locomotor suppression in rodents [6, 9, 10]. Dyn-fibers synapse onto DA terminals, and KOR is expressed on these same DA terminals as a heteroreceptor [11, 12]. This anatomical co-presence of Dyn/KOR structures on DA neurons suggests that the presynaptic DYN/KOR system modulates DA transmission. Indeed, KOR agonists decrease DA release and extracellular DA concentrations via the activation of KOR located on DA terminals, and the decreased dopaminergic neurotransmission is suggested to be involved in KOR agonist-induced CPA and locomotor suppression [10, 13–15]. Furthermore, recent studies have shown that the specific elimination of KOR in DA neurons abolishes KOR-agonist-induced place aversion and synaptic DA modulation, produces anxiolytic behavior and enhances cocaine sensitization [16, 17]. Mitogen activated protein kinase (p38 MAPK) in DA neurons and KOR in the dorsal raphe nucleus are involved in KOR mediated aversion [17, 18]. However, the downstream targets on which KOR triggered signaling cascades act to drive aversion and other behavioral actions are unknown.

The functional properties of DA transporter (DAT) on DA neurons are one of the critical factors in regulating synaptic DA for neurotransmission through clearance of released DA and behavior [19, 20]. It is well documented that several cellular protein kinases regulate DAT function via post-translational modifications such as phosphorylation [reviewed in and see references therein 21, 22, 23]. Activation or inactivation of intracellular kinase/phosphatase signaling cascades might occur through activation of homo- and hetero-receptors expressing on DA-neurons/terminals. Consequently, DAT mediated DA clearance and release can be regulated by receptors to modulate DA neurotransmission and behaviors. However, the role of DAT phosphorylation in a physiologically relevant model system has yet to be determined. Given the fact that KOR is expressed as a heteroreceptor on DAT expressing DA terminals [11, 12], using heterologous expression systems and native rat striatal tissue, we previously demonstrated that KOR agonists upregulate DAT function via ERK1/2 activation and that this regulation requires Thr53 phosphorylation of DAT (pT53-DAT) [24, 25]. However, a causative relationship of endogenous pT53-DAT to KOR-mediated DAT upregulation has not yet been investigated, and its contribution to KOR agonist-induced aversion and locomotor suppression is also unknown.

We recently used a knock-in mouse model (DAT-Ala53) carrying a non-phosphorylatable Ala at Thr53 phosphorylation site in DAT to examine the impact of in-vivo pT53-DAT on DAT function and animal behavior [26]. In the current study, we directly examined the causal role of in-vivo pT53-DAT in mediating KOR-linked DAT regulation, aversion-like behavior, and locomotor suppression using DAT-Ala53 and WT mice. We determined the effect of systemically administered KOR agonists on pT53-DAT levels along with DAT activity, kinetics and surface expression. We also investigated the genotype effects on locomotor suppression and CPA induced by KOR agonists. KOR agonists upregulated DAT activity by enhancing the total pT53-DAT, DAT-V_max_ and the surface levels of DAT and pT53-DAT in WT mice, but not in DAT-Ala53 mice. Furthermore, our results demonstrate that the lack of Thr53-DAT phosphorylation in DAT-Ala53 mice prevents locomotor suppression and CPA produced by KOR agonists. Nonetheless, DAT-Ala53 mice exhibit intact LiCl-induced CPA and morphine-induced hyperlocomotion and CPP. These results suggest that KOR-signaling mediated pT53-DAT exhibits a causal relation to dopaminergic neurotransmission and consequent behavioral outcomes such as locomotor suppression and aversion-like behavior caused by KOR agonism.

## Materials and methods

### Animals and housing

All animal studies and care were performed under the guidelines of the Virginia Commonwealth University Institutional Animal Care and Use Committee (IACUC), in accordance with the principles and procedures outlined in the National Research Council “Guide for the Care and Use of Laboratory Animals.” All mice were housed in a colony room (3-5 males per cage) and housed in a temperature and humidity-controlled environment under a 12 hr light/12 hr dark light schedule and were fed standard mouse chow (irradiated Teklad LM-485 diet) and autoclaved water. Our laboratory generated DAT-Ala53 knock-in mice on a C57BL/6J background, and their validation was thoroughly characterized and published in our previous study (Ragu Varman et al., 2021). 8-10 weeks wild type (WT) littermates and DAT-Ala53 homozygous male mice were used for behavioral and molecular experiments. All behavioral and biochemical experiments were conducted between 10 am and 5 pm.

### Experimental groups and drugs

Male WT or homozygous DAT-Ala53 mice were assigned to one of four treatment groups: saline + vehicle, saline + U69593 (0.1mg/kg, s.c.), nor-BNI (10 mg/kg, i.p.) + vehicle or nor-BNI + U69593 (where nor-BNI was given 24 hrs prior to vehicle or U69593). U69593 was dissolved in 20% propylene glycol in saline [27], and the vehicle served as 20% propylene glycol alone. Injectable grade isotonic saline (0.9% NaCl) solution was used to dissolve Nor-BNI, U50488 (10 mg/kg, i.p.), SalA (1mg/kg, i.p.), morphine (10 mg/kg, s.c. for locomotor assay, 3 mg/kg. s.c. for CPP) and LiCl (160 mg/kg, i.p.). All drugs were given in a volume of 10 µl/gm body weight. U69593 (Cat. No: U103), nor-binaltorphimine (nor-BNI; Cat. No: 5.01087), U50448 (Cat. No: D8040), morphine (Cat. No: M8777), LiCl (Cat. No: L-4408), and propylene glycol (Cat. No: P4347) were obtained from Sigma-Aldrich, Burlington, MA). Salvinorin A (SalA) was from RTI-11597-93-18, NIDA Drug Inventory Supply and Control System). The drug concentrations were selected according to previous studies that showed effective conditioning behaviors in mice [6, 28, 29].

### Preparation of synaptosomes and determination of DAT-mediated DA uptake and serotonin transporter (SERT)-mediated 5-HT uptake

WT and DAT-Ala53 male mice received vehicle or KOR agonist U69593 (0.1mg/kg, s.c.) and were returned to their home cage until experiment initiation. A separate group of animals received nor-BNI (10 mg/kg, i.p.) 24 hours before vehicle or KOR agonist administration. After 2 hours post-injection of vehicle or U69593, animals were decapitated, and quickly brains were removed and kept on ice. The dose of U69593 and preadministration time were selected based on the published study [27] that showed stimulation of DAT activity by U69593. Dorsal and ventral striatum were dissected on a cold dish and collected in 2 ml of ice-cold sucrose buffer (0.32 M sucrose in 5 mM HEPES, pH 7.4). The procedure for synaptosome preparation and DA and 5-HT uptake assay was adapted from our previous study [26, 30]. The tissue was homogenized using a Teflon-glass homogenizer, and the homogenized samples were centrifuged at 1000 g for 10 min at 4°C. After centrifugation, the supernatant was collected and centrifuged at 12,000 g for 20 min. The resulting pellet was resuspended in ice cold sucrose buffer. The protein concentration was quantified by a protein assay kit using BSA as a standard. Synaptosomes (30 μg of protein) were incubated in a total volume of 0.3 ml of KRH-assay buffer, pH 7.4 (120 mM NaCl, 4.7 mM KCl, 1.2 mM KH_2_PO_4_, 1.2 mM MgSO_4_ and 10 mM HEPES and 10 mM D-glucose) containing 0.1 mM ascorbic acid and 0.1mM pargyline. For DAT kinetic analysis, [^3^H] DA (20 nM) was mixed with unlabeled DA so that total DA concentration ranges from 25 nM to 1000 nM. Total DAT-mediated DA uptake (DAT-specific + nonspecific DA uptake) was determined by measuring DA uptake in the presence of NET-specific inhibitor nisoxetine (50 nM) to block NET-mediated DA uptake. To determine the nonspecific DA uptake, synaptosomes were preincubated with both the DAT and NET specific inhibitors (50 nM GBR-12909 and 50 nM nisoxetine respectively) at 37°C for 10 min followed by the addition of 10 nM [^3^H] DA (Cat No: NET673001MC; 18.32 Ci/mmol: Perkin Elmer, Santa Clara, CA). [^3^H] DA uptake was terminated at 5 min with the addition of GBR-12909 followed by rapid filtration over GF-B filters using Brandel Cell Harvester (Brandel Inc., Gaithersburg, MD). In parallel, SERT-mediated 5-HT uptake was determined. To determine the nonspecific 5-HT uptake, synaptosomes were preincubated with SERT-specific inhibitor (0.1 µM fluoxetine) at 37°C for 10 min. 5-HT uptake was initiated by the addition of 10 nM [^3^H] 5-HT (Cat No: NET498001MC; 24.82 Ci/mmol: Perkin Elmer, Santa Clara, CA). and the uptake was terminated at 5 min with the addition of fluoxetine followed by rapid filtration over GF-B filters. Filters were counted by liquid scintillation counter (MicroBeta2 LumiJET, PerkinElmer) to determine the radioactivity bound to the filters. Nonspecific DA uptake measured in the presence of GBR-12909 and nisoxetine was subtracted from the total [^3^H] DA uptake measured in the presence of NET inhibitor nisoxetine to determine the DAT-specific DA uptake. Nonspecific 5-HT uptake measured in the presence of fluoxetine was subtracted from the total [^3^H] 5-HT uptake to determine the SERT-specific 5-HT uptake.

### Surface protein biotinylation and determination of total and surface DAT and pT53-DAT levels

Surface protein biotinylation was carried out as described in our previous study [26]. Dorsal or ventral striatal synaptosomes from respective treatment groups and genotypes (under experimental groups and drugs) were used for surface protein biotinylation. Synaptosomes were incubated with the membrane-impermeable EZ link NHS-Sulfo-SS-biotin (Cat No: 21331, Pierce, Rockford, IL) (1 mg/1 mg protein) for 30 min at 4°C in ice-cold PBS/Ca/Mg buffer (138 mM NaCl, 2.7 mM KCl, 1.5 mM KH_2_PO_4_, 9.6 mM Na_2_HPO_4_, 1 mM MgCl_2_, 0.1 mM CaCl_2_, pH 7.3). At the end of the biotinylation and following centrifugation, the synaptosomes were suspended in the same buffer containing 100 mM glycine to quench excess NHS-Sulfo-SS-biotin. The biotinylated synaptosomes were pelleted by centrifugation and suspended in ice cold radioimmunoprecipitation assay (RIPA) buffer (10 mM Tris-HCl, pH 7.5, 150 mM NaCl, 1 mM EDTA, 1% Triton X-100, 0.1% SDS, and 1% sodium deoxycholate) supplemented with commercial cocktails of protease and phosphatase inhibitors [(1 µg/ml aprotinin, 1 µg/ml leupeptin, 1 µM pepstatin, and 250 µM phenylmethylsulfonyl fluoride; Cat. No: P8340, Sigma-Aldrich, Burlington, MA); (10 mM sodium fluoride, 50 mM sodium pyrophosphate, 5 mM sodium orthovanadate, and 1 µM okadaic acid (Cat. No: P0044, Sigma-Aldrich, Burlington, MA)]. The suspended synaptosomes were solubilized by passing through a 26-gauge needle six times followed by incubation at 4°C for 60 min and the lysate was collected by centrifugation at 25,000 g for 30 min. Biotinylated proteins were isolated by incubating the lysate/supernatant overnight with NeutrAvidin Agarose resin (Cat No: 29201, Pierce, Rockford, IL). The NeutrAvidin Agarose resin was washed three times with RIPA buffer by brief centrifugation and the biotinylated proteins (bound to the NeutrAvidin Agarose resin) were eluted by incubating the resin with 50 µl Laemmli sample buffer (62.5 mM Tris-HCl pH 6.8, 20 % glycerol, 2 % SDS and 5 % ß-mercaptoethanol). Proteins from total and bound fractions were separated by 7.5 % SDS-PAGE and transferred to polyvinylidene difluoride membranes (Bio-Rad, Hercules, CA). To reduce the background signal, the membrane blots were blocked by incubating with 5% BSA-TBS-Tween 20 buffer for 1 hour at room temperature. Total levels of DAT and pT53-DAT were determined using primary antibodies for DAT (mouse anti-dopamine transporter monoclonal, clone mAb16, Cat. No: MABN669, 1:2000 dilution, MilliporeSigma, Burlington, MA) and pT53-DAT (rabbit anti-phospho-Thr53 dopamine transporter polyclonal, Cat. No: p435-53, 1:2000 dilution, PhosphoSolutions Inc., Aurora, CO) respectively. After washing the membranes, corresponding species-specific horseradish peroxidase-conjugated secondary antibodies (1: 10000 dilution, Jackson ImmunoResearch Laboratories, Inc, West Grove, PA) were used to visualize immunoreactive bands by ECL reagents (Cat. No: GERPN2236, Amersham Biosciences, GE Healthcare, Pittsburgh, PA). The level of intracellular endoplasmic reticulum protein calnexin was determined by stripping the membranes and reprobing with anti-calnexin antibody (rabbit anti-calnexin polyclonal Cat. No: SPA 860, 1:2000, Enzo Life Sciences, Inc. Farmingdale, NY) to validate that only plasma membrane surface proteins were biotinylated and to ensure that equal protein was loaded in the gel and transferred to membrane. Expression levels of arbitrary protein band densities of pT53-DAT, DAT, and calnexin were quantified using NIH Image J software (version 1.48j). The protein bands were quantified by ensuring that results were within the linear range of the film exposure (HyBlot CL, Thomas Scientific, Swedesboro, NJ). The relative arbitrary protein band densities obtained for pT53-DAT and DAT in total fractions were normalized with calnexin.

### Locomotor activity measurement

Locomotor activity was measured in open field activity monitoring chambers (Med Associates, St. Albans, VT; Model ENV 510) in a soundproof box as described earlier [26]. The horizontal activity was recorded for 60 min and data were collected in 4 min bins as cm. WT and DAT-Ala53 mice were handled for 5 days before initiation of locomotor activity measurement. Mice received vehicle or U69593 alone or morphine alone or nor-BNI alone or nor-BNI + U69593 in a volume of 1 ml /kg and immediately placed in an activity monitor chamber, and activity was monitored for 60 min. Nor-BNI was administered 24 hours before vehicle or KOR agonist administration.

### Conditioned place aversion and preference study

Drug conditioning was performed using a three-compartment apparatus enclosed in a sound-attenuating cubicle (Med-Associates, St. Albans, VT, ENV3013). The apparatus consisted of white and black chambers (20 X 20 X 20 cm each), which were distinguished from one another by their different floor textures (white chamber with mesh and black chamber with rod: Med-Associates, ENV-3013WM and ENV-3013BR). The smaller grey intermediate compartment with a smooth PVC floor and doors leading to the black and white chambers served as a barrier between the place conditioning chambers. The compartments were equipped with photobeam strips, and the location, locomotion, and time spent by the subject in each compartment were recorded using the program provided by the manufacturer.

Mice were handled for 5 days before conditioning experiments. Conditioning KOR agonist U69593 was adapted as described [6, 16, 31]. On day 1, mice were given 15 minutes to explore both compartments for initial chamber preference (preconditioning test or pre-test), and the amount of time spent in each compartment was recorded. Mice were conditioned with U69593 (0.1mg/kg, s.c.) or vehicle on day 2 following the pre-test. For conditioning, mice were injected once daily for 6 days alternatively either with vehicle or U69593 and confined to a specific compartment for 15 min. Mice injected with U69593 were confined to the compartment where they spent more time during pre-test (preferred side) and mice injected with vehicle were confined to the compartment where they spent less time during pre-test (non-preferred side). nor-BNI was administered 24 hours before vehicle or KOR agonist administration. Vehicle control mice received vehicle daily for 6 days. Place conditioning preference was tested 24 hours after the last conditioning session in a drug-free state by allowing the animals free access to all three compartments for 15 minutes (post-conditioning test), and time spent in each compartment was recorded. Conditioned place aversion (CPA) to other KOR agonists U50448 and SalA, conditioned place preference (CPP) to µ-opioid receptor agonist morphine (morphine conditioning was done in non-preferred side) and CPA to non-opioid LiCl were determined using the same design as described above with the exception that there were six separate drug-conditioning sessions spread across three days, twice daily. After testing for initial chamber preference on day 1 (pretest), animals received saline in one chamber in the AM and U50448 or SalA or morphine or LiCl in the opposite chamber in the PM each day for 3 days (conditioning, 30 min duration). Post-conditioning test was conducted 24 hours after the last conditioning session in a drug-free state by allowing the animals free access to all three compartments for 15 min, and time spent in each compartment was recorded. The drug concentrations (given under *Experimental groups and drugs)* were selected according to previous studies that showed effective conditioning behaviors in mice [6, 28, 29]. Horizontal activity was recorded during each conditioning session to determine whether treatments affect activity. To determine conditioned preference, the time spent in the drug-paired side during the post-conditioning test was subtracted from their time during the pre-test. A drug is defined as aversive or producing CPA if the animal spends significantly less time in the drug-paired compartment than the vehicle-paired compartment during the test session. In contrast, a drug exhibits a rewarding effect (CPP) when the animal spends significantly more time in the drug-paired compartment than the vehicle-paired compartment during the test session.

### Statistical analysis

All values are expressed as Mean ± standard deviation (SD). Microsoft Excel (Mac-version 16.84) and GraphPad Prism 10 (RRID: SCR 002798 GraphPad, San Diego, CA) were used for data analysis, statistical evaluations, and to generate graphical representations. ImageJ 1.54g (National Institutes of Health, USA) was used to quantify bands from digitized films digitally. Figures were presented as bar graphs showing all points representing each subject or repeats. One-way or two-way analysis of variance was used, followed by post-hoc testing for pair-wise comparisons. Student’s t-tests were used to compare two datasets. A value of p < 0.05 was considered statistically significant. Specific statistical analyses, statistical data, and the significance of each experiment are included in the results section. In addition, comparisons between groups with p-values and the number of samples used in each experiment are reported in each figure legend.

## Results

### Acute KOR activation by systemic U69593 upregulates DAT activity, DAT-T53 phosphorylation, V_max_ and surface expression in WT mice, but not in DAT-Ala53 mice

Using in vitro cell culture model and ex vivo rat striatal synaptosomes we showed that KOR agonist SalA upregulates DAT function, V_max_, and surface expression in an ERK1/2-dependent manner [25]. In our subsequent study, we showed that ERK1/2 inhibition failed to inhibit/alter striatal DAT activity in DAT-Ala53 knock-in mice [26]. Therefore, it was essential to investigate how activation of KOR modulates DAT function and whether in-vivo KOR-mediated upregulation of endogenous DAT function requires DAT phosphorylation. Given the facts that (a) KOR-triggered ERK1/2 is mediating DAT upregulation, (b) DAT-T53 phosphorylation site is the signature canonical motif for ERK1/2, and (c) the absence of effects of ERK1/2 inhibition on DAT activity in DAT-Ala53 knock-in mice, we sought to investigate the role of DAT-T53 phosphorylation in KOR-mediated DAT upregulation in both ventral and dorsal striatum of DAT-Ala53 knock-in mice expressing normal DAT function and expression but lacking Thr53 phosphorylation (DAT-Ala53) by comparing with WT controls. KOR agonist, U69593 (0.1 mg/kg, s.c.) was systemically administered to DAT-Ala53 and WT control mice to activate KOR. DAT mediated DA uptake in ventral and dorsal striatum was determined using synaptosomes 2 hr U69593 post-administration. The dose, time, and route of administration were selected from previous publication demonstrating enhanced DAT activity by U69593 [27]. When compared to vehicle, U69593 administration significantly enhanced DAT-mediated DA uptake in both ventral and dorsal striatum of WT mice (ventral striatum: p = 0. 0003; dorsal striatum: p < 0.0001) but not in DAT-Ala53 mice (ventral striatum: p = 0. 9969; dorsal striatum: p = 0. 8526) (Fig. 1A & B). Two-way ANOVA showed significant effect of genotype (ventral striatum: F(1,4) = 59.35, p < 0.0015; dorsal striatum: F(1,4) =37.86, p < 0.0035) and treatment (ventral striatum: F(3,12) = 5.708, p < 0.0115; dorsal striatum: F(3,12) = 2.276, p < 0.1320) with significant genotype X treatment interaction (ventral striatum: F(3,12) = 7.860, p < 0.0036; dorsal striatum: F(3,12) = 13.84, p < 0.0003). To further confirm that KOR is involved in U69593-mediated upregulation of DAT function, we examined the effect of the long-acting KOR antagonist nor-BNI [32]. We injected KOR antagonist nor-BNI (10 mg/kg, i.p.) 24 hrs before U69593 administration. While nor-BNI alone did not affect the basal DAT function in WT, it prevented U69593-mediated upregulation of DAT function in both ventral (p < 0. 0018) and dorsal striatum (p < 0.0001). nor-BNI administration alone or with U69593 did not affect DAT function in DAT-Ala53 mice significantly (ventral striatum: p < 0. 1960 and dorsal striatum: p < 0.9811) (Fig. 1A & 1B). We also examined KOR agonist effect on SERT function in ventral and dorsal striatum of DAT-Ala53 to determine if T53 mutation in DAT affects SERT function. Consistent with published observation [33], systemic U69593 administration enhanced SERT-mediated 5-HT uptake in ventral and dorsal striatum of WT. Similar to WT, enhanced ventral and dorsal striatal SERT activity was observed in DAT-Ala53 mice following systemic U69593 administration (Fig. 1C & 1D). The two-way ANOVA revealed s significant effect of U69593 on ventral and dorsal striatal SERT activity in both WT and DAT-Ala53 mice (ventral striatum: F(1,4) =16.40, p < 0.016; dorsal striatum: F(1,4) = 9.25, p = 0.038) with no significant genotype effect (ventral striatum: F(1,4) = 0.90, p = 0.395; dorsal striatum: F(1,4) = 0.38, p = 0.567) or genotype X treatment interaction (ventral striatum: F(1,4) = 0.000, p = 0.927 and dorsal striatum: F(1,4) = 0.01=7, p = 0.895). These results indicate the specific effect of DAT-Ala53 mutation in KOR-mediated DAT regulation.

**Figure 1.**
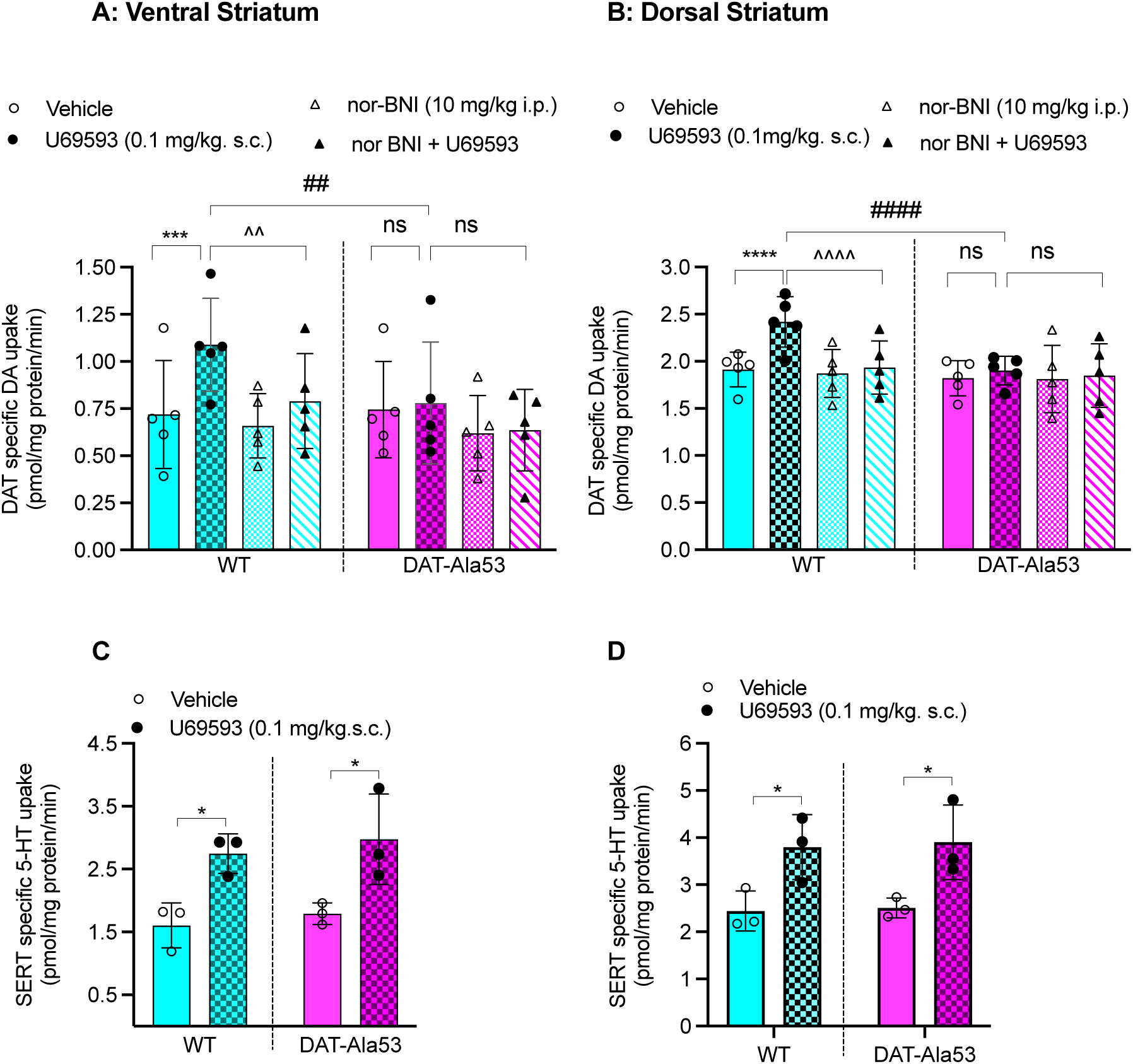
Effect of U69593 on DAT and SERT activities in ventral and dorsal striatal synaptosomes. Nor-BNI sensitive increase in DAT activity is observed in the ventral (A) and dorsal (B) striatal synaptosomes of WT, but not DAT-Ala3 mice following systemic U69593 administration. Drug administrations, synaptosome preparations, and DAT-specific ^3^H labeled DA uptake assays were performed as described in the materials and methods section. DAT-specific [^3^H] DA uptake is given in pmol/mg protein/ min and presented as Mean ± S.D. Each data point represents an individual mouse. When compared with vehicle control, U69593 increased DAT activity in the WT mice (ventral striatum: \*\*\**p* = 0.0003; dorsal striatum \*\*\*\**p* = 0.0001) but not in the DAT-Ala53 mice (ventral striatum: *p* = 0.997; dorsal striatum: *p* = 0.853). nor-BNI blocked U69593 effect on DAT activity in the WT mice (ventral striatum: ^^*p =* 0.002; dorsal striatum: ^^^^*p =* 0.0001). Significant difference between WT-U69593 and DAT-Ala53-U69593 (ventral striatum: ## *p =* 0.002; dorsal striatum: ####*p =*0.0001). ns: nonsignificant when compared between specific pairs as indicated in figures. WT (n = 5) and DAT-Ala53 (n = 5) mice. Data were analyzed by two-way RM ANOVA with Tukey’s multiple comparisons. U69593 increased SERT activity in ventral (C) and dorsal (D) striatal synaptosomes of WT (ventral striatum: \**p =* 0.026; dorsal striatum: \**p =* 0.042) and DAT-Ala53 (ventral striatum: ^*p =* 0.022; dorsal striatum: \**p =* 0.037) mice. Data were analyzed by two-way ANOVA with Sidak’s multiple comparisons.

To examine the kinetic properties of DAT-mediated DA uptake, saturation analysis was performed in synaptosomes obtained from ventral and dorsal striatum of WT and DAT-Ala53 mice administered with U69593 or vehicle. Consistent with our previous study [26], there was no effect of genotype on the V_max_ and K_m_ of DAT mediated DA uptake in ventral and dorsal striatum of vehicle administrated mice (Fig. 2). Compared with vehicle administration, U69593 administration in WT mice increased DAT V_max_ in both ventral and dorsal striatum (ventral striatum-± SD, vehicle: 16.08 ± 0.3625 pmol/mg/min, U69593: 23.85 ± 1.117 pmol/mg/min and dorsal striatum-± SD, vehicle: 19.983 ± 0.886 pmol/mg/min, U69593: 24.403 ± 1.651 pmol/mg/min) without altering K_m_ (ventral striatum-± SD, vehicle: 168.93 ± 18.45 nM, U69593: 157.16 ± 6.017 nM and dorsal striatum-± SD, vehicle: 122.78 ± 43.927 nM, U69593: 109.69 ± 16.919 nM) (Fig. 2). However, in DAT-Ala53 mice, compared with vehicle control, systemic U69593 did not produce significant change in DAT V_max_ in both ventral and dorsal striatum (ventral striatum-± SD, vehicle: 15.22 ± 0.931 pmol/mg/min, U69593: 16.20 ± 1.491 pmol/mg/min and dorsal striatum-± SD, vehicle: 18.62 ± 0.869 pmol/mg/min, U69593: 19.73 ± 0.899 pmol/mg/min). K_m_ was unaltered in ventral and dorsal striatum (ventral striatum-± SD, vehicle: 145.36 ± 21.553 nM, U69593: 165.76 ± 14.164 nM; p = 0.767 and dorsal striatum-± SD, vehicle: 128.10 ± 21.287 nM, U69593: 128.80 ± 30.153 nM; p = 0.639) (Fig. 2). Two-way ANOVA of V_max_ showed significant effect of genotype (ventral striatum: F(1,2) = 93.10, p < 0.011; dorsal striatum: F(1,4) = 12.2, p < 0.025) and treatment (ventral striatum: F(1,2) = 90.0, p < 0.011; dorsal striatum: F(1,4) = 25.9, p < 0.007) with significant genotype X treatment interaction (ventral striatum: F(1,2) = 21.2, p < 0.011 and dorsal striatum: F(1,4) = 72.9=7, p < 0.001). These results suggest that the phosphorylation of Thr53 is required to enhance endogenous DAT activity in ventral and dorsal striatum following KOR activation.

**Figure 2.**
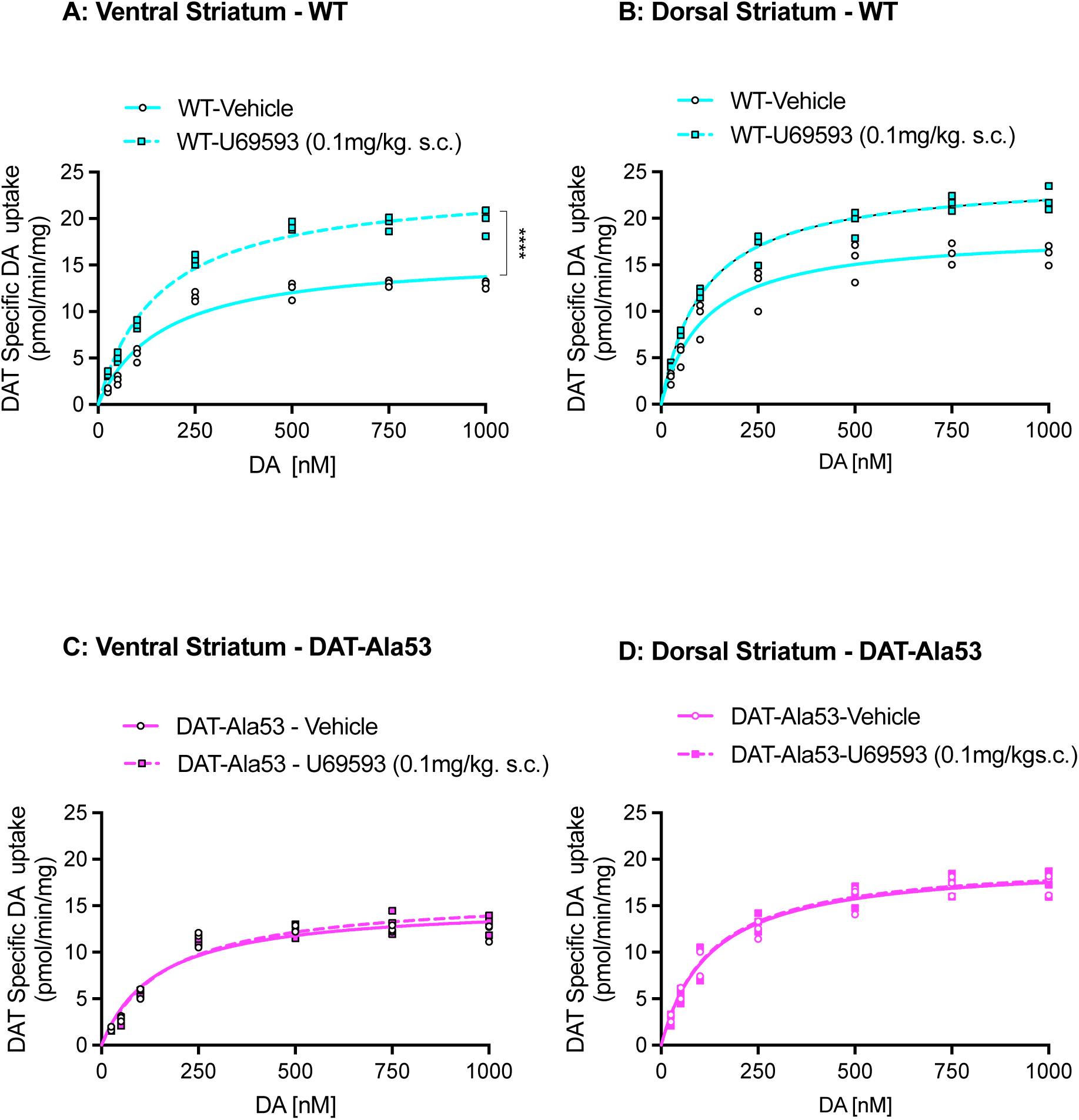
U69593-induced increase in DAT V_max_ is blunted in the ventral and dorsal striatum of DAT-Ala53 mice. Drug administrations, synaptosome preparations, and DAT-specific DA transport kinetic analysis were performed as described in the materials and methods section. Maximal velocity (V_max_) and substrate (DA) apparent affinity (K_m_) of DAT were determined by using Michaelis-Menten equation (Prism). DA uptake values (pmol/mg protein/min) are presented as Mean ± S.D. WT (n = 3) and DAT-Ala53 (n = 3). When compared with vehicle, U69593 increased DAT V_max_ significantly in (A) ventral striatum (\*\*\*\**p* = 0.0001) and not significantly in (B) dorsal striatum (*p* = 0.999) of WT mice. U69593 failed to increase DAT V_max_ in (C) ventral (*p* = 0.572) and (D) dorsal striatum (*p* = 0.999) of DAT-Ala53 mice. U69593 did not alter K_m_ in ventral and dorsal striatum of both WT and DAT-Ala53 mice. Data were analyzed using two-way ANOVA with Bonferroni’s multiple comparisons.

To study the mechanism underlying the elevated DAT function following U69593 administration, we sought to determine whether systemic U69593 mediated upregulation of DAT activity and V_max_ is accompanied by increased pT53-DAT and cell surface DAT. Membrane surface protein biotinylation and immunoblotting with specific antibodies to pT53-DAT, total DAT (DAT), and calnexin were used to quantify total and surface resident of DAT, pT53-DAT and calnexin in ventral and dorsal striatum of WT and DAT-Ala53 mice (Fig. 3). Systemic administration of U69593 in WT mice significantly increased pT53-DAT without altering total DAT and intracellular marker calnexin in ventral and dorsal striatum (Fig. 3A, C, E). Consistent with increased DA uptake and V_max_, systemic U69593 administration significantly increased surface (biotinylated) levels of DAT and pT53-DAT in ventral and dorsal striatum of WT mice (Fig. 3B, D, F). In contrast to WT, systemic U69593 administration failed to alter surface DAT level in the ventral and dorsal striatum of DAT-Ala53 mice (Fig. 3B, F) without changes in total DAT (Fig. 3A, E). Two-way ANOVA revealed significant genotype effect (ventral striatum: pT53-DAT-F(1,4) = 176.2, p = 0.0002; DAT-F(1,4) = 12.70, p = 0.023) and dorsal striatum: pT53-DAT-F(1,4) 661.7, p < 0.0001; DAT-F(1,4) = 20.08, p = 0.011); and treatment effect (ventral striatum: pT53-DAT-F(1,4) =13.43, p = 02153; DAT-F(1,4) = 33.24, p < 0.005 and dorsal striatum: pT53-DAT-F(1,4) = 46.69, p = 0.0024; DAT-F(1,4) =29.53, p < 0.006) with significant genotype X treatment interaction in both ventral and dorsal striatum (ventral striatum: pT53-DAT-F(1,4) = 13.18, p = 0.0221; DAT-F(1,4) = 25.95, p = 0.007 and dorsal striatum: pT53-DAT-F(1,4) = 46.61, p = 0.0024; DAT-F(1,4) =17.66, p = 0.014). As reported in our previous publication [26], the pT53-DAT antibody did not detect pT53-DAT apart from some non-specific bands in ventral and dorsal striatum of DAT-Ala53 mice. Notably, the intracellular endoplasmic reticulum marker calnexin is undetectable in biotinylated fractions, indicating that synaptosomes were intact and intracellular DAT protein was not biotinylated and recovered along with surface biotinylated DAT or pT53-DAT (Fig. 3B). Thus, the observed biotinylated DAT represents resident DAT on the surface plasma membrane. These findings demonstrate that in-vivo DAT-T53 phosphorylation in the ventral and dorsal striatum is required for KOR-mediated upregulation of DAT-specific DA transport by enhancing the surface density of DAT.

**Figure 3.**
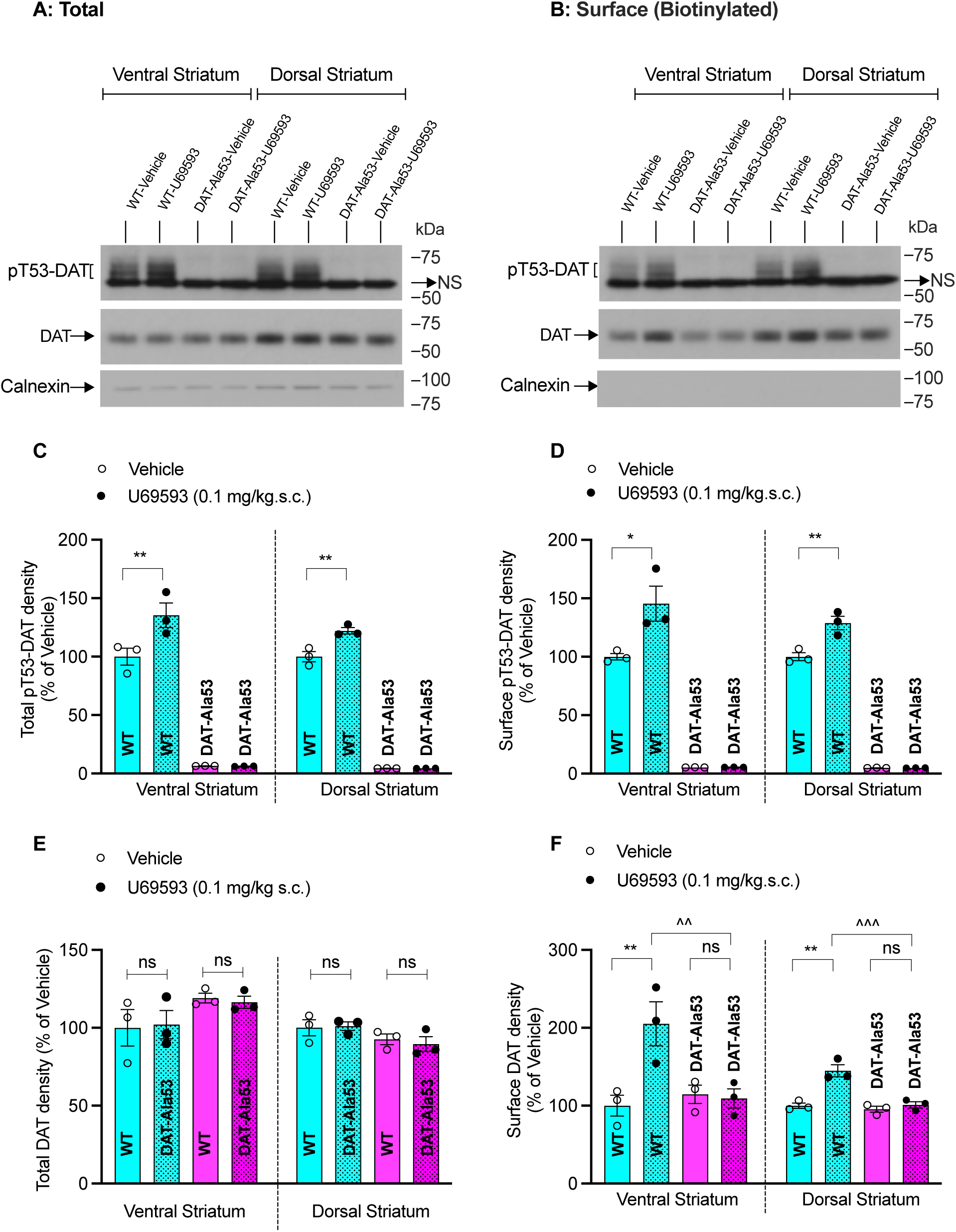
U69593 enhances total and surface DAT-Thr53 phosphorylation and surface DAT expression in ventral and dorsal striatum of WT but not the DAT-Ala53 mice. Drug administrations and synaptosome preparations are given under the materials and methods section. Using indicated specific antibodies, total and surface expression of pT53-DAT, DAT and calnexin were visualized and quantified following surface protein biotinylation. Representative immunoblots from three independent experiments show changes in (A) total and (B) surface (biotinylated) pT53-DAT (∼60-55kDa), DAT (∼65 kDa) and calnexin (∼90 kDa) in the ventral and dorsal striatum of WT and DAT-Ala53 mice. The specificities of DAT and pT53-DAT immunoreactive bands by the antibodies used were established in our previous publication [26]. Bar graph shows quantified arbitrary band densities of total pT53-DAT (C), DAT (E) and biotinylated surface pT53-DAT (D), and DAT (F). Each data point represents an individual mouse, and the data presented as Mean ± SEM. WT (n = 3) and DAT-Ala53 (n = 3) mice. Compared with vehicle control, U69593 increased total and surface pT53-DAT levels and surface DAT expression without altering the total expression of DAT and calnexin in the ventral and dorsal striatum of WT but not DAT-Ala53 mice. \**P* = 0.05, \*\**P* < 0.01, ^^ *P* < 0.01, ^^^ *P* < 0.001, ns: nonsignificant. Comparisons between specific pairs are indicated in the figures. Data were analyzed by two-way ANOVA with Bonferroni’s multiple comparisons.

### KOR activation by U69593 does not elicit locomotor suppression in DAT-Ala53 knock-in mice

It is known that systemic administration of KOR agonists in rodents induces reduced locomotor activity [6, 10]. Since KOR activation reduces DA release [10, 13–15, 34], and our results showed that KOR activation also increases DA clearance through DAT-T53 phosphorylation-dependent trafficking mechanism, we hypothesized that DAT-T53 phosphorylation may have a causal relationship to the behavioral effects of KOR agonists. To test our hypothesis, we compared the effect of systemic U69593 on the modulation of locomotor activity in WT and DAT-Ala53 mice. We selected U69593 dose (1.0 mg/kg) that is shown to affect locomotor activity [6]. Consistent with published studies [6], when compared to vehicle administration, systemic U69593 administration significantly decreased locomotor activity (total distance traveled in 60 min) of WT mice (p < 0.0001) (Fig. 4). However, U69593 administration did not have any significant effect on the locomotor activity of DAT-Ala53 mice (p > 0. 1357) (Fig. 4). Locomotor activity measured in 5 min bins is shown in (Fig. 4C, D). Two-way ANOVA analysis of the total activity for the duration of 60 min showed significant effect of genotype F(1,14) = 13.94, p < 0.0022) and U69593 administration (F(1.14) = 12.70, p < 0.0031) with significant genotype X U69593 interaction (F(1,14) = 8.482, p < 0.0114). Pretreatment with long-acting KOR antagonist, nor-BIN attenuated U69593-induced locomotor suppression in WT mice (p < 0. 0009) suggesting the involvement of KOR activation. Moreover, the significant difference between WT and DAT-Ala53 mice with respect to the effect of U69593 on locomotor activity (p < 0.0006) suggests an essential role of DAT-T53 phosphorylation in KOR-mediated locomotor suppression (Fig. 4).

**Figure 4.**
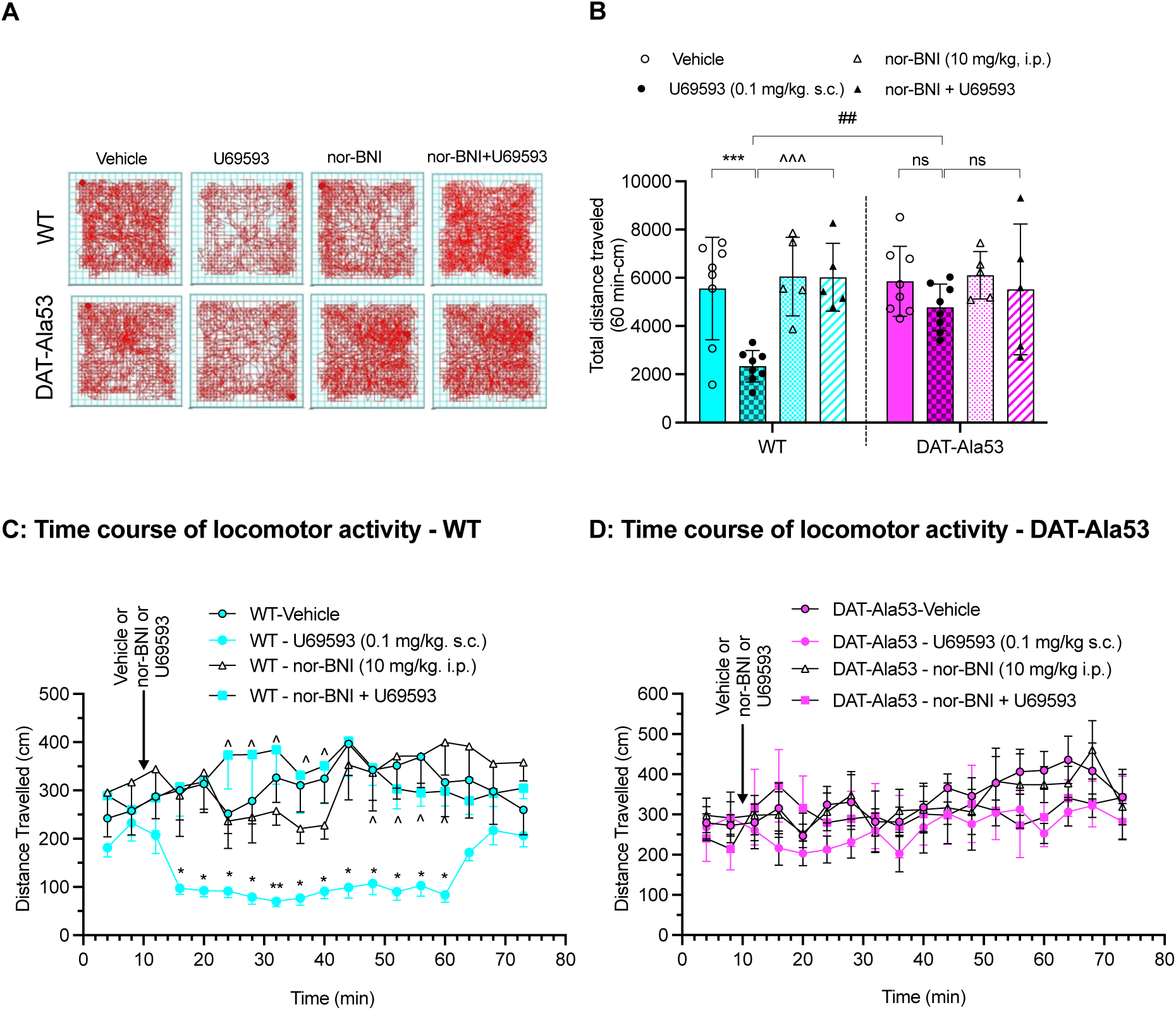
U69593-induced suppression of locomotor activity is blunted in DAT-Ala53 mice. Representative activity traces of 60 min locomotor activity (A), total distance traveled for 60 min (B) and time course of locomotor activity measured in 4 min bins over 60 min period (C & D) post administration of vehicle or U69593 or nor BNI or pre-nor-BNI following U6953 in WT and DAT-Ala53 mice are shown. Systemic administration of vehicle or U69593 and locomotor monitoring were performed as described under the materials and methods section. Each data point represents an individual mouse, and the data is presented as Mean ± SEM. WT (n = 6) and DAT-Ala53 (n = 6) mice. Comparison of U69593 effects on locomotor activity between genotypes revealed that U69593 decreased locomotor activity in WT (\*\*\**p* = 0.001) but not in DAT-Ala53 (*p* = 0.999) mice. nor-BNI blocked U69593 induced locomotor suppressing effect in WT (^^^*p* = 0.001). ##*p* = 0.0001 WT-U69593 versus DAT-Ala53-U69593. ns: nonsignificant (*P* = 0.999). Comparisons between specific pairs are indicated in the figures. Data were analyzed by two-way ANOVA with Bonferroni’s multiple comparisons.

### The KOR agonists do not induce CPA in DAT-Ala53 knock-in mice

It is well known that KOR agonists produce CPA in rodents [6, 9, 10, 16, 31, 35, 36]. To investigate whether DAT-T53 phosphorylation-dependent DAT regulation plays a role in KOR agonist-triggered CPA, we tested the effect of systemic administration of KOR agonists (U69593, U50488 and SalA) on WT and DAT-Ala53 mice (Fig. 5). To keep continuity and comparisons with published literature, the dose of KOR agonists and conditioning procedures were adapted from published CPA protocols [6, 9, 10, 16, 31, 35, 36] and as described under Materials and Methods. As expected, all three agonists tested produced CPA in WT mice. Multiple comparisons indicate that WT mice administered with U69593 or U50488 or SalA showed significant CPA (U69593: p < 0. 0001; U50488: p = 0. 0268; SalA: p = 0.017) compared to vehicle administration (Fig. 5A, C). Pretreatment with long-acting KOR antagonist, nor-BNI blocked U69593 triggered CPA (p = 0. 0001) suggesting the involvement of KOR activation (Fig. 5A). Interestingly, U69593 and U50488 did not induce significant CPA in DAT-Ala53 mice (U69593: p = 0. 404; U50488: p = 0. 268). On the other hand, SalA elicited significant CPA in DAT-Ala53 mice (p = 0. 001) (Fig. 5C). However, the SalA-induced CPA was significantly lower in DAT-Ala53 mice compared to WT mice (p = 0.01). There was significant genotype difference between WT and DAT-Ala53 mice in response to KOR agonists (U69593: p = 0. 0013; U50488: p = 0. 0032, SalA: p = 0.01). Two-way mixed ANOVA showed significant effect of treatments (U69593: F(1,31) = 25.67, p < 0.0001; U50488: F(1,7) = 18.2, p = 0.004; SalA: F(1,7) = 7.363, p = 0.030) and significant differences in the CPA produced by all three KOR agonists between genotypes (U69593: F(1,31) = 8.389, p < 0.01; U50488: F(1,7) =18.2 p = 0.004; SalA: F(1,7) = 20.60, p = 0.003) with significant genotype X treatment interaction except for SalA (U69593: F(1,31) = 9.044, p < 0.001; U50488: F(1,7) = 9.36, p < 0.018; SalA: F(1,7) = 2.793, p = 0.139). Furthermore, there were no significant differences in post-test movement counts between genotypes following treatment with all three KOR agonists (Fig. 5B, D). These observations indicate that KOR agonists cause aversion-like behavior by modulating KOR-mediated DAT regulation through DAT-T53 phosphorylation.

**Figure 5.**
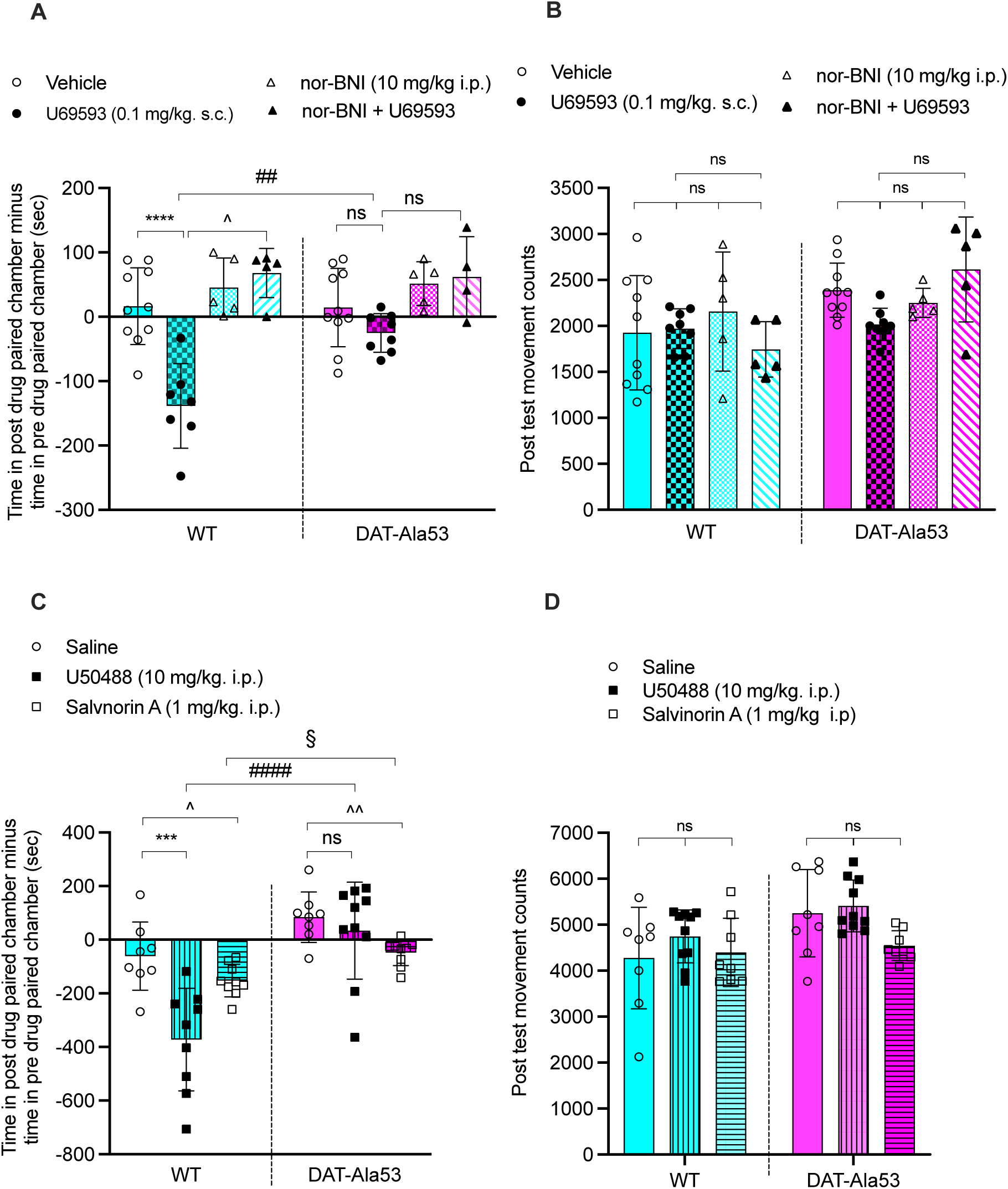
U69593, U50488 and SalA-induced conditioned place aversion are blunted in DAT-Ala53 mice. Mice were conditioned with indicated KOR agonists as described under the materials and methods section. Each data point represents an individual mouse, and bars are plotted as average values ± S.D. The post-conditioning effects of vehicle and U69593 (A) and the post-test movement counts (B) are shown. The number of mice used for vehicle (n=10) and U69593 (n=8) for each genotype. U69593 produced significant CPA in WT (\*\*\*\**p* < 0.0001) but not in DAT-A53 (ns = 0.453) mice in comparison with corresponding vehicle controls. A significant difference in the CPA effect of U69593 between WT and DAT-Ala53 mice (##*p* = 0.0013). nor-BNI blocked U69593-induced CPA in WT mice (^*p* = 0.025). The post-conditioning effects of U50488 and SalA (C) and the post-test movement counts (D) are presented. U50488 and SalA produced significant CPA in WT (\*\*\**p* < 0.0003; ^*p* = 0.045 respectively). DAT-A53 mice displayed no CPA to U50488 (ns = 0.999) and attenuated CPA to SalA (§*p* = 0.015). ####*p* = WT-U50488 *versus* DAT-Ala53-U50488. ^^*p* = 0.006 DAT-Ala53-saline *versus* DAT-Ala53-SalA. ns: nonsignificant compared between specific pairs as indicated in figures. The number of mice used WT-saline (n=8), WT-U50488 (n=9), WT-SalA (n=9), and DAT-Ala53-saline (n=8); DAT-Ala53-U50488 (n=10) and DAT-Ala53-SalA (n=9). Data were analyzed by Mixed-effects analysis with Bonferroni’s multiple comparisons.

### Morphine-induced enhancement of locomotion is not affected in DAT-Ala53 knock-in mice

We examined the effect of morphine, a mu-opioid receptor agonist, on the modulation of locomotor activity to ensure that the lack of locomotor suppression in DAT-A53 mice following U69593 administration is not a generalized phenomenon. As shown in Fig. 6, systemic morphine administration (10 mg/kg, s/c.) triggered locomotor activity in both WT and DAT-Ala53 mice. There were significant morphine treatment effect (F(1,9) = 37.62, p = 0.0002) in both genotypes without genotype X treatment interaction (F(1,9) = 5.084, p = 0.0506). We observed that DAT-Ala53 mice display higher morphine-induced locomotor activity (p = 0.024) compared to WT mice. These results indicate that the absence of U69593 mediated locomotor suppression in DAT-Ala53 mice is not a generalized phenomenon.

**Figure 6.**
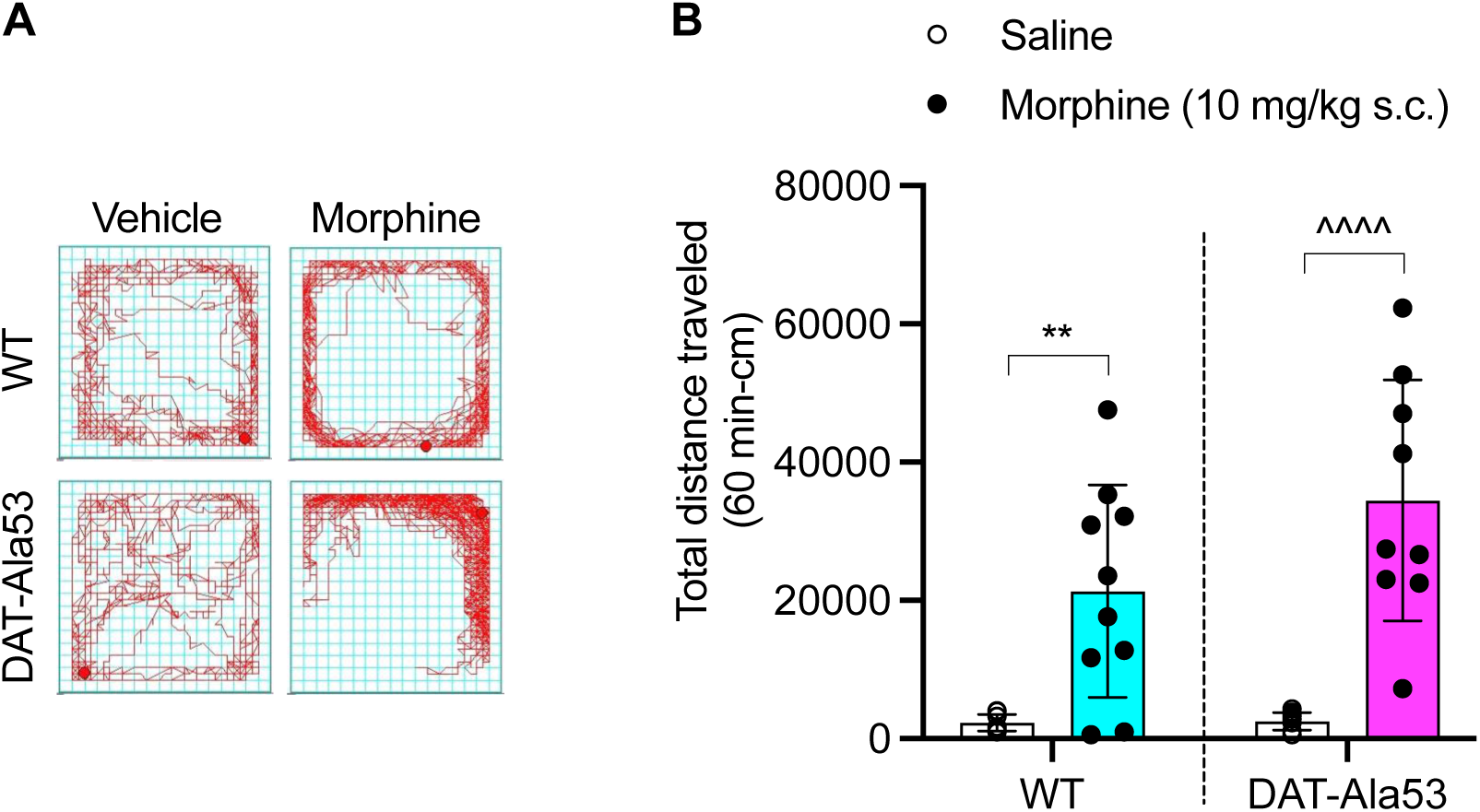
Morphine-triggered locomotor activity is similar in WT and DAT-Ala53 mice. Administration of morphine and its effect on locomotor activity were described in the materials and methods section and the data presented as mean ± S.D. WT (n = 10) and DAT-Ala53 (n = 9). Representative activity traces of 60 min locomotor activity (A) and total distance traveled for 60 min (B) post administration of vehicle or morphine in WT and DAT-Ala53 mice are shown. Bonferroni’s multiple comparative analysis of the impact of morphine on locomotor activity showed that morphine evoked significant locomotor activation in WT (\*\**p* = 0.003) and DAT-Ala53 (^^^^*p* = 0.0001) mice. Data were analyzed by Mixed-effects analysis with Tukey’s multiple comparisons.

### Morphine-induced CPP and LiCl-induced CPA are intact in DAT-Ala53 knock-in mice

To examine further whether the attenuated aversive effects of KOR agonists observed in DAT-A53 mice (Fig. 5) occur by affecting learning and performance of the conditioned response or by producing generalized malaise, we examined the conditioned response of morphine and non-opioid lithium chloride (LiCl). As presented in Fig. 7A, and consistent with published literature [29, 37], morphine (3 mg/kg, s.c.) conditioning produced place preference in WT mice. Similar to WT mice, DAT-Ala53 mice also exhibited morphine-induced CPP. Post-test movement counts did not differ between genotypes (p = 0.089, t=1.28, df = 14) (Fig. 7B). Two-way ANOVA indicated significant morphine conditioning effect in both genotypes (F(1,26) = 35.60, p = 0.0001) and no significant genotype effect (F(1,26) = 5.28, p = 0.474) with no significant genotype X treatment interaction (F(1,26) =0.757, p = 0.392). We used the same cohort of WT and DAT-Ala53 mice used for KOR agonists triggered conditioning behavior after two weeks of washout period to examine the conditioned aversive effects of another nonopioid drug, lithium chloride. As predicted [28, 38], LiCl (160 mg/kg, i.p.) produced conditioned aversion in WT (Fig. 7C). Surprisingly, the conditioned aversive effect of lithium chloride was intact in DAT-Ala53 mice (Fig. 7C). Post test movement counts did not differ between genotypes (p = 0.648, t = 0.466, df = 14) (Fig. 7D). Two-way ANOVA indicated significant LiCl conditioning effect (F(1.28) = 28.70, p = 0.0001) in both genotypes without significant genotype X treatment interaction (F(1,28) = 0.0101, p = 0.9206). Overall, intact morphine CPP and LiCl CPA observed in DAT-Ala53 mice as seen in WT mice indicate that the lack of KOR agonist-induced aversive behavior in DAT-Ala53 mice is not due to altered learning and/or performance of conditioned response.

**Figure 7.**
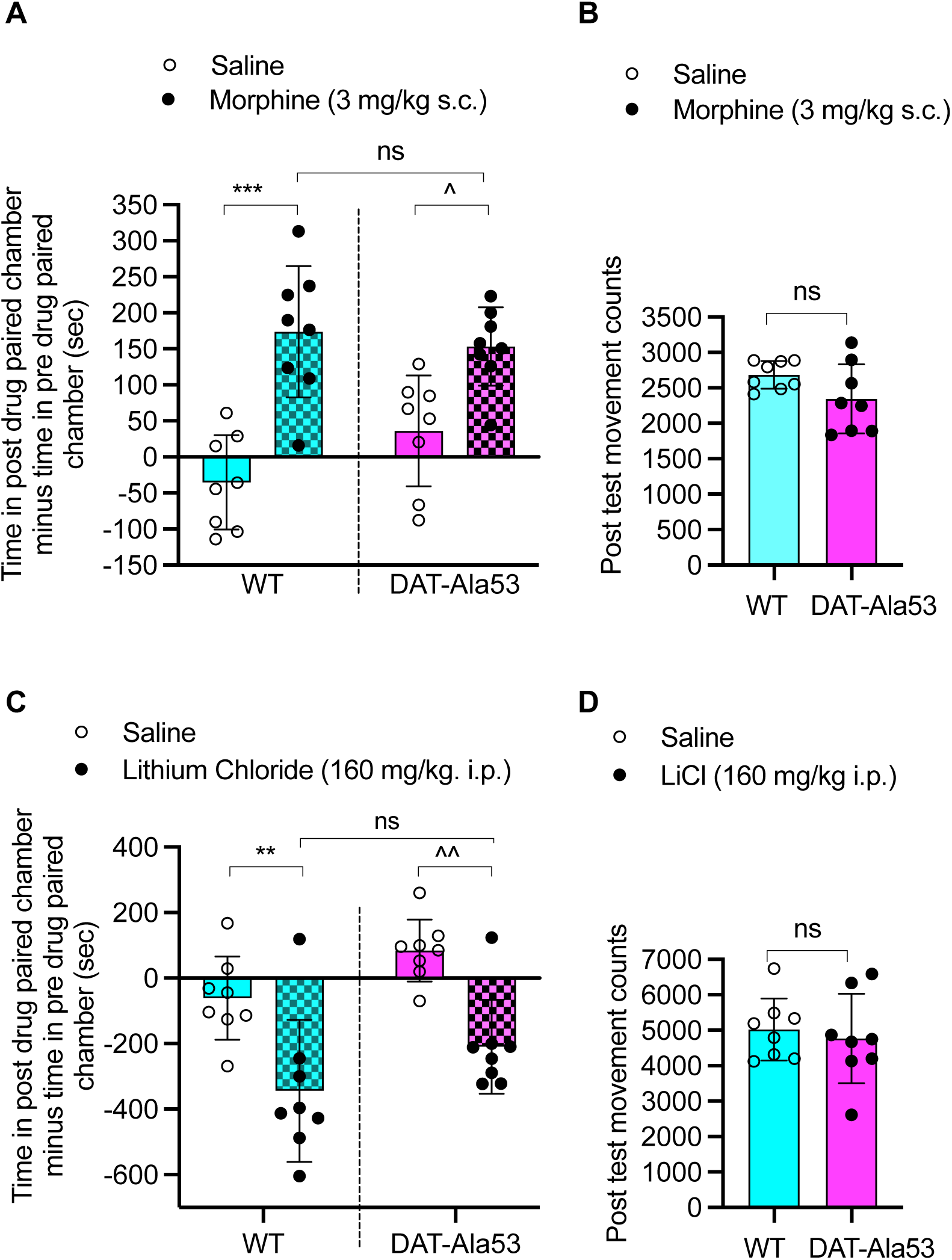
DAT-Ala53 mice display similar morphine-CPP and LiCl-CPA as that of WT. The same cohorts of WT and DAT-Ala53 mice used above in Fig. 5 & 6 for measuring locomotor activity were used to condition with morphine or LiCl or saline, and their effects on place preference or aversion were determined as described under materials and methods. The post-test of conditioned effects of morphine (A), LiCl (C), and total locomotor distance traveled during post-test (morphine (B) & LiCl (D) in comparison with WT and DAT-Ala53 are presented. Each data point represents an individual mouse, and bars are plotted as averages ± S.D. Conditioning with morphine produced significant CPP in both WT (\*\*\*\**p* = 0.0001) and DAT-Ala53 (^*p* = 0.017) mice and showed no differences in morphine CPP between WT and DAT-Ala53 mice (ns = 0.943) WT (n = 8) and DAT-Ala53 (n = 8). LiCl conditioning produced significant CPA in both WT (\*\**p =* 0.0047) and DAT-Ala53 (^^*p =* 0.0032) mice and showed no difference between WT and DAT-Ala53 (ns *=* 0.309). The number of mice used WT-saline (8), WT-LiCl (8), and DAT-Ala53-saline (8) and DAT-LiCl (8). Comparison of between specific pairs as indicated in figures. Data were analyzed by two-way ANOVA with Tukey’s multiple comparisons.

## Discussion

The primary findings from the current study reveal that KOR-mediated DAT-T53 phosphorylation is a primary causal post-translational mechanism underlying the effects of KOR agonists on locomotor suppression and aversion-like behaviors in male mice. The current data establish that in-vivo DAT-T53 phosphorylation on DA-terminals is required for (a) KOR-mediated upregulation of DAT functional kinetics and surface expression, (b) KOR agonist-mediated locomotor suppression and (c) KOR agonist-mediated aversive-like behaviors.

KOR is expressed on DA terminals, and its effects of decreasing DA release have been implicated in the aversive and locomotor suppressive properties of KOR agonists [7, 10, 13, 14]. Supportive of this notion, mice lacking KOR in DA neurons exhibit abolished KOR agonist-triggered aversion and decreased extracellular DA levels [16]. Furthermore, re-expressing KOR in DA neurons of KOR knock-out mice recovered KOR agonist-induced aversion [17]. These studies indicate that KOR expression and activation in DA neurons are essential for KOR-mediated decreased extracellular DA levels and aversive and locomotor suppressive behavior in mice.

Extracellular DA levels available for subsequent neurotransmission are dynamically governed by synchronized regulation of DA release and uptake. Functional DAT activity on DA-terminals governs extracellular DA dynamics [19]. The fact that DAT kinetic properties and surface expression are dynamically regulated by kinase-mediated phosphorylation and that KOR activation triggers DAT-T53 phosphorylation to stimulate DAT function and surface expression in a heterologous cell culture model [24, 25] reveals a potential role of in-vivo DAT-T53 phosphorylation in KOR-mediated DA clearance and KOR-induced aversive behavior and locomotor suppression. However, an investigation of this hypothesis is lacking to date. To test this hypothesis, in the current study, we investigated KOR-agonist mediated DAT regulation, aversion, and locomotor suppression using DAT-Ala53 knock-in mouse model lacking T53-DAT phosphorylation but exhibiting unaltered DAT functional expression [26].

Systemic administration of KOR agonist, U69593 to WT mice stimulates DAT-mediated DA uptake in both ventral and dorsal striatum in a nor-BNI-sensitive manner. The increased DAT activity is associated with higher transport velocity (V_max_) and unaltered DAT affinity to DA (K_m_). Additionally, U69593 triggers pT53-DAT and increases surface DAT and pT53-DAT levels in the WT. In the absence of the DAT-T53 phosphorylation site in DAT-Ala53 mice, systemic U69593 failed to trigger DAT activity, V_max_ and surface expression suggesting that KOR-mediated in-vivo DAT upregulation is dependent on the phosphorylation of DAT at Thr53 site, which is also the ERK1/2 site [26, 39].

KOR agonists cause locomotor suppression in mice, which is absent in KOR-KO mice [6, 10]. Indeed, systemic KOR agonist U69593 suppressed the spontaneous open-field activity in WT mice, an effect blocked by KOR-antagonist nor-BNI pretreatment. This confirms that activation of KOR is involved in suppressing locomotor activity by U69593. Remarkably, systemic U69593 failed to alter locomotor activity in DAT-Ala53 mice. However, µ-opioid receptor agonist morphine administration stimulated locomotor activity in DAT-Ala53 mice similar to that of WT mice, suggesting the specific role of pT53-DAT in the effect of KOR agonist. Since a parallel decrease in extracellular DA and locomotor suppression is observed following KOR agonism, diminished dopaminergic transmission has been implicated in KOR agonist effects on locomotor activity [10]. DAT plays a critical role in regulating extracellular DA and locomotor activity [19]. Our findings that the absence of KOR-mediated DAT functional regulation and KOR-agonist-induced locomotor suppression in DAT-Ala53 mice suggest that the phosphorylation of DAT at T53 mediated by KOR signaling modulates DA clearance and, in conjunction with the evidence that KOR signaling/activation also decreases DA release [40] consequently, may cause locomotor suppression.

Extensive studies revealed that KOR agonists produce CPA in rodents [6, 9, 10, 16, 31, 35, 36] which requires KOR expression in DA neurons [16, 17]. In agreement with previous findings, systemic administration of three KOR agonists, U69593, U50488, and SalA produced CPA in WT mice. KOR antagonist, nor-BNI prevented U69593-induced CPA suggesting the specific involvement of KOR in KOR agonist-induced CPA. Surprisingly, U69593 and U50488 completely failed to elicit CPA and SalA produced significantly attenuated CPA in DAT-Ala53 mice, suggesting the need for DAT-T53 phosphorylation for KOR agonists to induce CPA. Deficiencies in learning and performance of the conditioned response in DAT-Ala53 mice would arise without KOR agonist-mediated CPA. We tested and compared the conditioned response to non-opioid LiCl and µ-opioid receptor agonist morphine in WT and DAT-Ala53 mice. It is known that LiCl produces CPA [38] and morphine induces CPP [29, 37]. Our data show that LiCl-induced CPA and morphine-induced CPP are intact in DAT-Ala53 mice as seen in WT, suggesting that DAT-Ala53 mice are able to learn and respond to conditioned drugs. Thus, the absence of KOR agonist-mediated CPA was not due to any defect in the learning and performance of the conditioned response.

It has been reported that KOR-triggered p38 MAPK activation via GRK3/arrestin in ventral tegmental area-DA neurons is required for KOR-mediated CPA in mice [17]. However, one study demonstrated that ß-arrestin-2 knock-out mice exhibit KOR-agonist-induced CPA similar to its WT littermates/counterparts [6]. Activation of KOR in dorsal raphe nucleus (DRN) by stress stimuli, downstream GRK3-dependent activation of p38 MAPK and serotonin transporter (SERT) stimulation in the ventral striatal serotonergic terminals are required to elicit KOR-mediated CPA [18, 33]. Moreover, it has been demonstrated that SERT is required for KOR-mediated CPA [33]. While these studies demonstrated the possible involvement of both DA and 5-HT signals in KOR-mediated behaviors, an interaction between these two systems as primary causal relationship to the complex KOR-induced CPA is yet to be investigated. KOR-mediated activation of p38 MAPK may regulate functional aspects of several substrates within DA or 5-HT neurons via phosphorylation, and the nature of these substrates is unknown. KOR-mediated upregulation of SERT activity in the ventral and dorsal striatum is evident in DAT-Ala53 mice. The fact that KOR agonists failed to elicit CPA in these mice suggests that KOR-p38 MAPK mediated SERT upregulation is not independently engaging in aversive properties of KOR activation but probably integrates DA transmission via DAT-Thr53 phosphorylation. However, the possible compensatory changes due to constitutive DAT-Thr53Ala mutation contributing to KOR agonist-induced CPA and locomotor suppression cannot be ruled out. Identifying such neuronal substrate due to constitutive DAT-Thr53Ala mutation will provide further insights into the downstream effector of KOR/ERK1/2 mediated DAT-T53 phosphorylation. Importantly, to our knowledge, the present study identifies DAT-T53 phosphorylation as the primary causal mechanism in KOR-induced CPA for the first time. Since (a) ERK1/2 inhibition failed to modulate DAT function in DAT- Ala53 mice [26], (b) KOR agonist-mediated DAT upregulation is ERK1/2-dependent [25] and (c) DAT-T53 is the ERK1/2 phosphorylation site [39], we interpret that KOR agonists upregulate DAT function at DA terminals through ERK1/2-dependent DAT-T53 phosphorylation eliciting CPA and locomotor suppression.

Additionally, understanding the source of DYN and potential circuits that lead to KOR activation in DA terminals and cell soma will illuminate brain region-specific role of DAT-T53 phosphorylation [41]. It has been thoroughly established that stress triggers endogenous dynorphin release, activates KOR signaling, and promotes aversive and pro-addictive effects [8, 42–44]. It would be informative to investigate if stress stimuli mediate aversive effects via DYN/KOR-DAT- T53 phosphorylation. The current study used only male animals, and KOR-mediated DAT regulation and aversive and locomotor suppressive behaviors of KOR agonists in female WT and DAT-Ala53 mice remain to be determined. It is known that female rodents display differential DA release, ERK1/2 activation, DAT-T53 phosphorylation, and behavioral effects following KOR agonisms [45–51]. Further investigation is required to understand the dimorphic role of sex in DYN/KOR-DAT-T53 phosphorylation signaling that translates to behavioral outcomes.

KOR activation triggers Gαi-mediated signaling followed by GRK3/ß-arrestin 2-dependent downstream signaling cascades [52, 53]. It has been hypothesized that GRK3/ßarrestin 2- dependent p38 MAPK activation contributes to KOR-induced aversion and sedation in mice, while Gαi-mediated signaling is attributed to antinociceptive and antipruritic effects [42]. Based on these concepts, G-protein-biased KOR agonists are developed with the promise to have therapeutic potential without causing adverse effects such as aversion [reviewed in and see references therein 54, 55]. However, specific neuronal substrates that target KOR signaling specific to Gαi- and GRK3/ß-arrestin 2-dependent signaling are largely unknown. For instance, a synthetic analog of SalA RB-64 is a G-protein-biased KOR agonist and exhibits antinociception without affecting locomotor suppression. However, it should be noted that RB-64 induces CPA similar to unbiased KOR agonists SalA and U69593 in WT and ß-arrestin-2 knock-out mice. These observations suggest multiple unknown KOR intracellular signals and their potential causative substrate(s) contributing to the behavioral effects of biased and unbiased KOR agonists. Present study demonstrates that KOR-mediated DAT regulation, locomotor suppression, and aversive behaviors are absent in DAT-Ala53 mice, and thus, we conclude that DAT-T53 phosphorylation is a prerequisite molecular link in the adverse effects of KOR agonism.

## Conclusion

It is well known that KOR agonists exhibit adverse effects, such as sedation, aversion, dysphoria, depression, and enhance rewarding effects of psychostimulants [reviewed in and see references therein 43, 56, 57-59]. However, the specific KOR signaling pathway and its downstream molecular link of neuronal targets driving the adverse properties of KOR agonists are unclear. Our studies showed that KOR-mediated DAT-T53 phosphorylation is a primary causal post-translational mechanism underlying the effects of KOR agonists on locomotor suppression and aversion-like behaviors. The current results suggest that the DAT-T53 phosphorylation site or other DAT-regulatory motifs are potential therapeutic targets to fine-tune DA neurotransmission and form a basis for developing effective pharmacotherapeutics that minimize the adverse effects of KOR agonism.

## Acknowledgments

This work was supported by the National Institute of Health (RO1 DA054694) to SR and LDJ.

